# Evaluating anthropogenic noise impacts on animals in natural areas

**DOI:** 10.1101/171728

**Authors:** Alexander C. Keyel, Sarah E. Reed, Kathryn Nuessly, Elizeth Cinto-Mejia, Jesse R. Barber, George Wittemyer

## Abstract

1. Noise pollution is detrimental to a diversity of animal species and degrades natural areas, raising concern over the expanding footprint of anthropogenic noise on ecosystems. To guide management of noise sources, modeling tools have been developed to quantify noise levels across landscapes.
2. We demonstrate how to model anthropogenic noise using sound propagation models, including noise from point, line, and polygon sources. In addition, we demonstrate three ways of evaluating spatially-explicit noise impacts, by identifying where noise 1) exceeds a sound level threshold, 2) is audible, or 3) has the potential to mask species communications. Finally, we examine approaches to mitigate these noise impacts on animal species.
3. Noise sources in locations more favorable to sound propagation (e.g., locations with long, unobstructed lines-of-sight) will have a disproportionate impact on the surrounding area. We demonstrate how propagation models can identify sites with smaller acoustic footprints or sites that would benefit from additional noise-control measures.
4. Modeling decisions, such as choice of sound propagation model, sound source information, and the quality of the input data, strongly influence the accuracy of model predictions. These decisions can be guided by comparing model predictions to empirical data when it is available.
5. **Synthesis and applications**: Here, we demonstrate an approach for modeling and assessing anthropogenic noise sources across a landscape. Our versatile approach allows refining propagation outputs for species-specific questions as well as the quantitative evaluation of management alternatives. While the results are presented in the context of particular species, these approaches can be applied more generally to a wide range of taxa or used for multispecies assessments.

## Introduction

Anthropogenic noise affects species’ occupancy, behavior, distribution, reproduction, physiology, and ultimately fitness (Shannon *et al*. 2015). Noise can be an invisible source of habitat degradation (Ware et al. 2015), influence trophic interactions (e.g., predator-prey dynamics, Francis, Ortega & Cruz 2009), and change the provision of ecosystem services (Francis *et al*. 2012). Although most noise studies have focused on birds, terrestrial noise has been shown to affect a wide variety of taxa, including mammals, reptiles, amphibians, and invertebrates (Bowles *et al*. 1999; Morley, Jones & Radford 2014; Shannon *et al*. 2015). Consequently, there is increasing interest in characterizing and mitigating the impacts of noise pollution on biodiversity (e.g., Mullet *et al*. 2016).

With increased awareness of the threats posed to ecological systems by noise, several approaches to model noise propagation across landscapes have been developed (e.g., Kragh *et al*. 2002; Ikelheimer & Plotkin 2005; Reed, Boggs & Mann 2012; Keyel *et al*. 2017). Sound propagation models provide a means of assessing current and predicted noise levels and evaluating noise propagation under alternative management options (Harrison, Clark & Stankey 1980; Reed, Boggs & Mann 2012) or future scenarios (Dumyahn & Pijanowski 2011). As such, the application of propagation modeling can provide rapid and cost-effective insights for planning or management decisions to mitigate potential noise impacts on animals (e.g., management of snowmobile noise in Yellowstone National Park, Jacobson 2013). Here, we demonstrate approaches to implement sound propagation models for three applied case studies of the effects of anthropogenic noise on animal species.

We focus our analysis on energy-related development and motorized recreation, two noise sources that can substantially increase sound levels in natural areas (e.g., Harrison, Clark & Stankey 1980; Ramirez Jr & Mosley 2015). Noise from natural gas extraction has been shown to reduce species’ occupancy from large areas of habitat, interfere with species’ hunting behavior, alter species’ physiology, and influence trophic interactions (Bradshaw, Boutin & Hebert 1998; Bayne, Habib & Boutin 2008; Francis *et al*. 2011; Blickley *et al*. 2012; Mason, McClure & Barber 2016). Recently, noise from gas compressors was shown to be more important than canopy cover in explaining habitat selection in secondary cavity-nesting birds (Kleist *et al*. 2017). Recreational noise, too, has been shown to directly, negatively affect species’ behavior (Brattstrom & Bondello 1983; Karp & Root 2009), and noise is hypothesized to be an important factor driving the negative effect of motorized recreation on species (Harrison, Clark & Stankey 1980; but see Reimers, Eftestøl & Colman 2003). While studies directly examining noise effects from motorized recreation are lacking, recreational activities are identified as a threat to 23–26% of species listed under the United States Endangered Species Act, particularly from motorized, off-road vehicle use (i.e., all-terrain vehicles, four-wheel drive vehicles, motorcycles, snowmobiles, and dune buggies, Losos *et al*. 1995). A recent review of recreational impacts found that ~45% of studies of summer-season motorized recreation and ~80% of snow-based, winter motorized recreation had negative effects on species (Larson *et al*. 2016).

In this manuscript, we demonstrate the steps and highlight key considerations for modeling sound levels from point, line, and area sound sources. In addition, we demonstrate three approaches to estimate potential impacts of noise propagation on animal species: 1) where noise exceeds a threshold (Shannon *et al*. 2015), 2) where it is audible (e.g., ISO 389-7), 3) and where it has the potential to interfere with intraspecific communication (Lohr et al. 2003). Finally, we discuss important considerations when developing and applying sound propagation model outputs to management questions.

## Materials and methods

### OVERVIEW

We introduce our general modeling framework in the context of three case studies where the noise production occurred from a point (natural gas extraction), a linear feature (trail-based recreation), and a defined area (designated area-based recreation). We used Sound Mapping Tools V4.4 (SMT, Keyel *et al*. 2017, Keyel et al *in review*) with ArcGIS (10.3, 10.4, ESRI, Redlands, CA) to evaluate the acoustic impacts using publicly-available data sets (see Table 1, code used to run the analyses given in Appendix 1).

**Table 1.** Data used for the three case studies.

| Variable | Shale Ridge | Bangs Canyon | Stanislaus National Forest |
| --- | --- | --- | --- |
| Nearest weather station | Grand Junction | Grand Junction | South Lake Tahoe<sup>7</sup> |
| Weather station WBAN | 23066<sup>1</sup> | 23066<sup>1</sup> | 93230<sup>1</sup> |
| Year used for weather conditions | 2014 | 2014 | 2014 |
| Months used for weather conditions | September | July | January |
| Mean temperature (°C) | 19.8 | 26.4 | 2.5 |
| Mean relative humidity (%) | 44.3<sup>2</sup> | 29.7<sup>2</sup> | 50.0<sup>2</sup> |
| Mean wind speed (kph) | 12.0 | 13.6 | 5.5 |
| Modal wind direction (°) | 120 | 45 | 30 |
| Land Cover | LANDFIRE<sup>3,4</sup> | LANDFIRE<sup>3,4</sup> | SNOW |
| Elevation | NED<sup>4,5</sup> | NED<sup>4,5</sup> | NED<sup>4,5</sup> |
| Source data | Drill rig<sup>6</sup>, gas compressor<sup>6</sup> | All-terrain vehicle<sup>6</sup> | Snowmobiles<sup>6</sup> |
| 1/3 octave band range | 125-2000 Hz | 125-2000 Hz | 2500 Hz |

<sup>1</sup> Weather data acquired from QCLCD files (NOAA 2015).
<sup>2</sup> Relative humidity was calculated using the August-Roche-Magnus approximation (Alduchov & Eskridge 1996; McNoldy 2017), and a single average value was computed for the focal month.
<sup>3</sup> (LANDFIRE 2012).
<sup>4</sup> Resampled to 30 × 30 m cell size and converted to the appropriate UTM zone.
<sup>5</sup> National Elevation Data, 1 arc-second resolution, (USGS 2013).
<sup>6</sup> Drill rig measurements made by E. Brown, reported in Keyel *et al.* (2017), gas compressor measurements made by J. Barber, reported in Keyel *et al. (in review)*, all-terrain vehicle source data from Harrison *et al.* (1980), snowmobile data provided by D. Joyce, National Park Service Natural Sounds and Night Skies Division.
<sup>7</sup> The closest weather station was Bridgeport Sonora Junction (WBAN 433) but this had no data for the year and month of interest.

SMT provides an easy-to-use ArcGIS interface for several existing sound models, SPreAD-GIS (Harrison, Clark & Stankey 1980; Reed, Boggs & Mann 2012), NMSIMGIS (Ikelheimer & Plotkin 2005; Keyel *et al*. 2017), and a GIS implementation of ISO 9613-2 (ISO 9613-2). These models have previously been used to address natural resource-related questions (e.g., Sunder 2003; Barber *et al*. 2011). As the models require point inputs, lines and polygons were represented by points. While each case study uses a different focal species, the methods outlined here are applicable to a wide range of taxa. All dB values reported here are A-weighted Sound Pressure Levels re: 20 μPa (dBA) unless otherwise noted (i.e. case study 3); one-third octave band ranges used in the weightings are given in Table 1.

### CASE STUDY 1: GAS DEVELOPMENT IN SHALE RIDGES MANAGEMENT AREA, CO

To demonstrate application of the models to point sources, we examined a subset of potential wells planned for natural gas extraction through the Shale Ridges Master Leasing Plan in the Shale Ridges Management Area (39.3 N 108.3 W; BLM 2015a). We examined the expected increase in noise levels on land designated as a Bureau of Land Management (BLM) Area of Critical Environmental Concern (ACEC) due to drilling and from a hypothetical on-site gas compressor station at four proposed well sites. We used the SPreAD-GIS model (Reed, Boggs & Mann 2012) from Sound Mapping Tools to make spatially-explicit sound level predictions for each well location for the drilling and operation phases (see Table 1 for data sources). We assessed where mule deer (*Odocoileus hemionus*) might be displaced within the ACEC by gas exploration activities using a threshold-based approach. Mule deer were chosen due to their sensitivity to natural gas development (Sawyer *et al*. 2006; Sawyer, Kauffman & Nielson 2009; Northrup, Anderson & Wittemyer 2015; Johnson *et al*. 2016). We used a 45 dBA 1 s L_eq_ threshold as the level at which mule deer would be displaced, empirically estimated for a proxy species, caribou (*Rangifer tarandus caribou*; see Appendix 2 for derivation; Bradshaw et al. Bradshaw, Boutin & Hebert 1997), as hearing among ungulates is similar (Heffner & Heffner 2010). Finally, we repeated the procedure with a systematic grid of points spaced 500 m apart to evaluate the potential impact of alternative well placement locations.

### CASE STUDY 2: MOTORIZED RECREATION IN BANGS CANYON, CO

Bangs Canyon (38.93 N 108.5 W), adjacent to Colorado National Monument and located near Grand Junction, CO, is managed by the BLM for motorized recreation, non-motorized recreation, and wildlife. We examined where motorized recreation (represented by an all-terrain vehicle, ATV) could be audible to humans above natural background sound levels for one part of the management area. Humans were selected as the indicator species because of the detailed understanding of human audibility (ISO 389-7), and humans are a useful proxy for many species given that our hearing is similar to or better than that of many wild animals (e.g. see audiograms in Fay 1988; Buxton *et al*. 2017). Audibility was calculated with reference to background levels given by Harrison et al. (1980). Noise levels and background levels were used to calculate a cumulative d’ statistic based on human sensitivity for each one-third octave frequency band (ISO 389-7). The ATV was considered audible any time 10^∗^log_10_(d’) reached or exceeded an empirically-derived threshold of 7.3 (Fidell, Pearsons & Sneddon 1994). To conduct the analysis, trails within 200 m of highways were excluded from the analysis based on the assumption that the highway would be the dominant source of noise in these areas. The motorized route was broken into a series of points to simulate a single ATV traveling at ~6 m s^−1^ sampled every 20 seconds, which resulted in an approximately 120 m point spacing along the line. The SPreAD-GIS model was run for each motorized point using source levels reported by Harrison et al. (1980), and we considered the audibility impact in two ways. First, we looked at where on the non-motorized trails an ATV would be audible by calculating audibility based on a single ATV present at all of the points along the motorized trail. Second, we examined which locations along the motorized trail were most responsible for this impact on the non-motorized trail, to prioritize any mitigation measures or development of alternative routes. To accomplish this, the length of non-motorized trail where each motorized point was audible was computed (length was approximated by examining the area affected for a 1 m wide trail).

### CASE STUDY 3: SNOWMOBILES IN STANISLAUS NATIONAL FOREST, CA

The USDA Forest Service has proposed a 3471 ha snowmobile recreation area within the Stanislaus National Forest, CA, USA (38.514 N, 119.92 W). Here, we examined the potential for snowmobile noise to mask avian communications within a focal area encompassing a portion of the open recreation area, derived from the Stanislaus National Forest Land and Resource Management Plan (USDA Forest Service 2010), and its surroundings. The impact of the snowmobile area (polygon) was assessed using a systematic grid of points spaced 30 m apart. We modeled standard and next-generation four-stroke snowmobiles, running at a speed of 13.4 m s^−1^ (48.3 km h^−1^, source data provided by D. Joyce, National Park Service). Model predictions were first made for a single snowmobile of each type at every grid point. Then, the maximum sound level from any point was used to evaluate the potential impact of the snowmobile area. Finally, the maximum sound level results for a single snowmobile were compared to the results from eight snowmobiles by scaling the single snowmobile results by a factor of eight.

We tested the potential for snowmobiles to mask the peak frequency of White-breasted Nuthatch (*Sitta carolinensis*) vocalizations by evaluating the noise levels within the corresponding one-third octave band (2500 Hz, Nelson 2015a; b). Peak frequency of nuthatch vocalizations was extracted using Raven Pro (Bioacoustics Research Program 2014). We used the NMSIMGIS model (Ikelheimer & Plotkin 2005) from Sound Mapping Tools due to its ability to model snow-covered ground and its greater frequency range than SPreAD-GIS.

## Results

In the first case study considering natural gas development in the Shale Ridges Management Area, the four proposed wells were predicted to raise sound levels above 45 dBA for 0, 11.6, 69.1, 76.3 ha during drilling and 0, 6.9, 39.0, 43.0 ha during operation of a compressor station (Fig. 1, for wells 1, 2, 3, 4, respectively). When alternative locations were considered, potential acoustic impacts varied across the landscape, with wells 3 and 4 among the locations with the greatest possible acoustic impact. Selection of alternative locations within 1 km could substantially reduce impacts (Fig. 2).

**Fig. 1.**
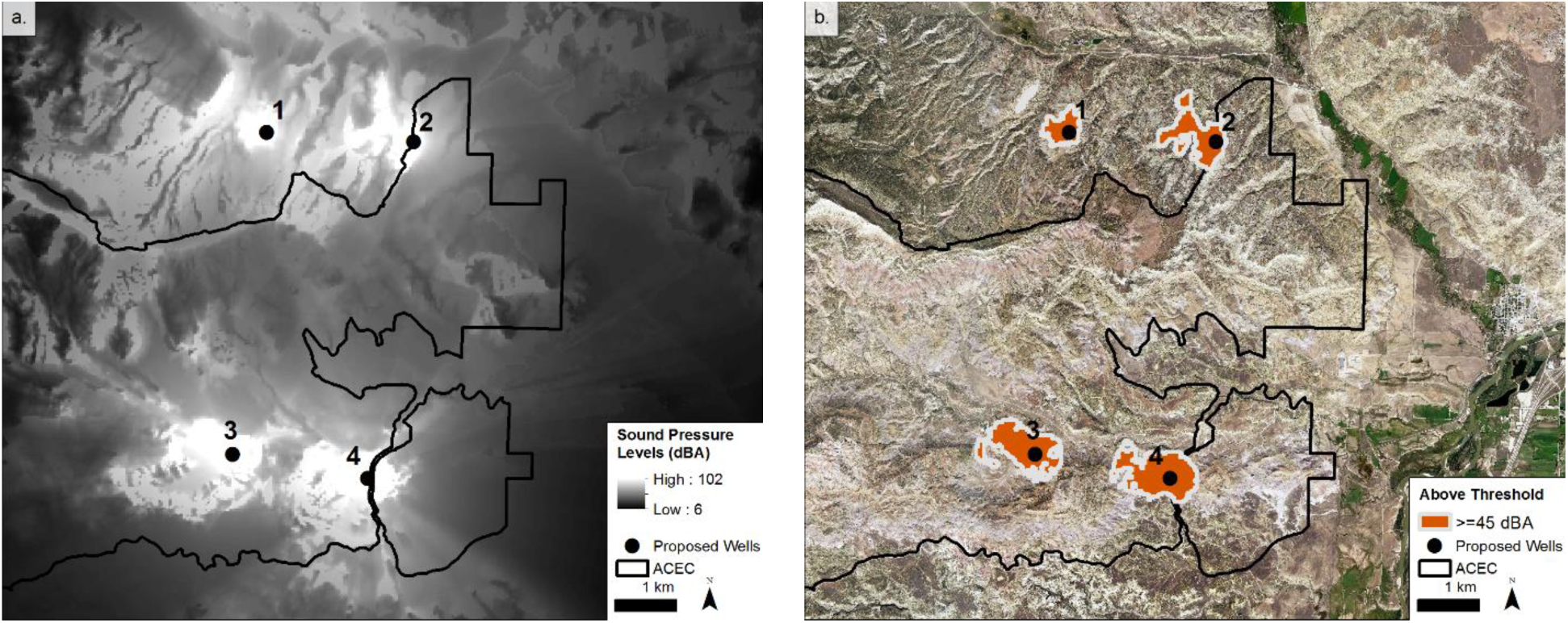
The predicted acoustic impact of drilling four new wells (1 - 4). (a) The predicted sound pressure levels of drilling the well sites are displayed, while in (b) only the areas where sound pressure levels would meet or exceed 45 dBA are shown. The Area of Critical Environmental Concern (ACEC) is outlined in black.

**Fig. 2.**
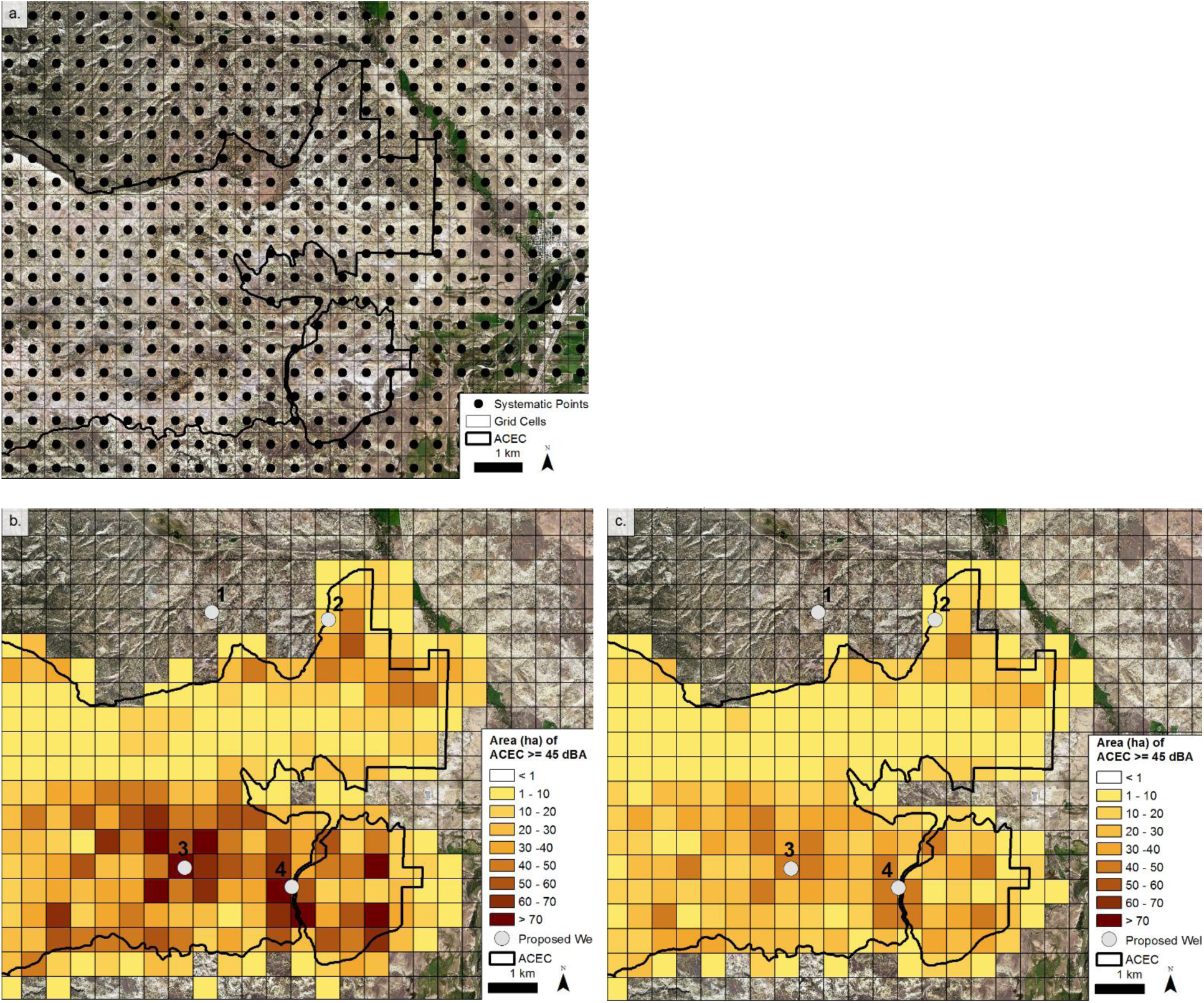
The potential impact on mule deer within the Area of Critical Environmental Concern (ACEC) was evaluated for (a) systematic points across the landscape. The impact of each systematic point was extrapolated to the entire grid cell, and each cell was color-coded according to the area of the ACEC that would be elevated above 45 dBA during (b) drilling and (c) by operation of a hypothetical on-site compressor station at that point. Note that the actual spatial extent above a 45 dBA threshold of each systematic point (as was shown in Fig. 1b) is not shown, rather the color coding provides an index to the spatial extent that would be affected by drilling or compressor operation at that point (the darker the shading the greater the area affected). The locations (but not the sound levels) of the four proposed well sites (1–4) are included for context.

In the second case study considering summer motorized recreation in Bangs Canyon, ATVs were predicted to be audible over 16% of the non-motorized trails (Fig. 3a). For most locations along the non-motorized trails, a single ATV would be inaudible due primarily to the barrier effects of intervening terrain and distance from the motorized trail (Fig. 3a). As in the first case study, not all points along the motorized trail were equal in their acoustic impacts, with seven point locations having a much larger acoustic impact on the non-motorized trail than the others (Fig. 3b).

**Fig. 3.**
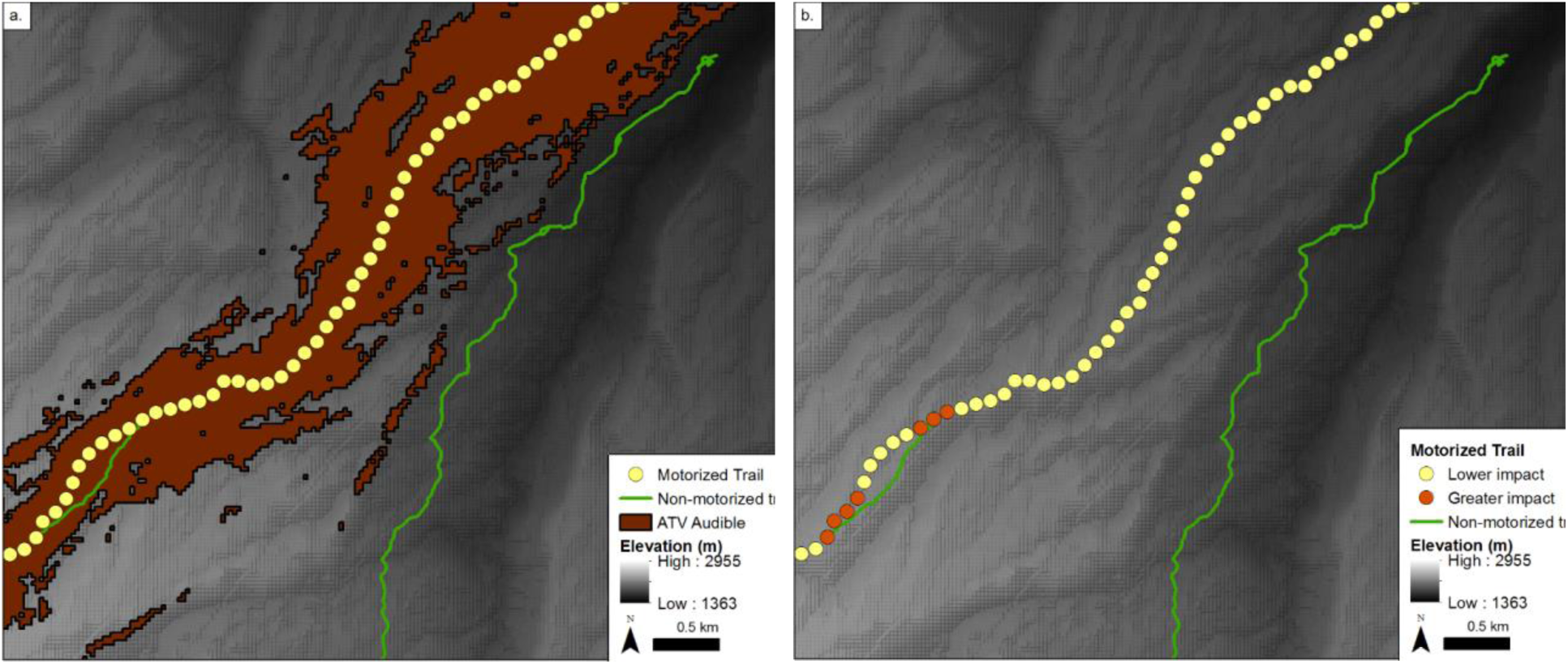
(a) Audibility of a single ATV traveling a motorized vehicle trail in Bangs Canyon, CO. Audibility was defined based on human hearing abilities (ISO 389-7), and was defined as a cumulative d’ statistic at or above 7. (b) The relative impact of individual sections of the motorized trail on the non-motorized trail highlight potential targets for management action. Lower impact points affected < 1 m of the non-motorized trail, while greater impact points affected ≥ 1 m. Elevation is from a hillshaded 1 arc-second digital elevation model (USGS 2013).

In the third case study considering winter motorized recreation in Stanislaus National Forest, snowmobiles differed in their potential to mask White-breasted Nuthatch vocalizations (Fig. 4). The next-generation snowmobiles produced lower sound levels in the 2500 Hz one-third octave band used by White-breasted Nuthatches.

**Fig. 4.**
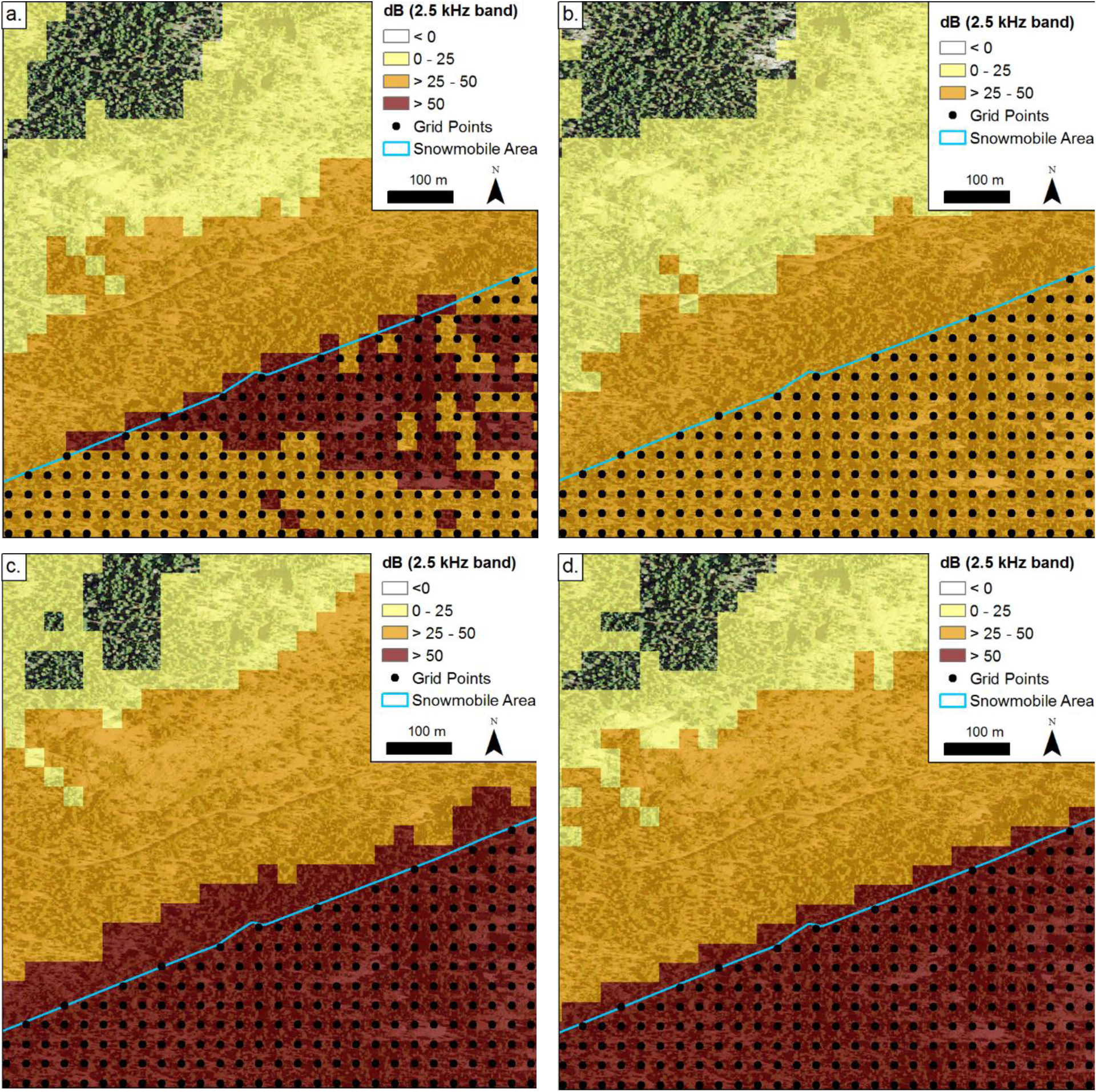
The potential for snowmobiles to mask White-breasted Nuthatch vocalizations in the 2500 Hz band differed by snowmobile type and number. Standard four-stroke snowmobiles (a, c) produce more acoustic energy in the 2500 Hz one-third octave frequency band than snowmobiles utilizing next-generation technology (b, c). Increasing the number of snowmobiles from one (a, b) to eight (c, d) raises the maximum sound level within the snowmobile area to above 50 dB for both snowmobile types (aerial photo from USDA-FSA-APFO 2016).

## Discussion

A spatially-explicit prediction of noise propagation using Sound Mapping Tools allowed the quick and inexpensive assessment of potential noise impacts on animals and evaluation of alternative management scenarios. Three case studies revealed spatial patterns of potential impacts that are much more complex than estimates based on distance alone. The first case study demonstrated the important role topography can play in sound propagation (Embleton 1996). Well locations differed substantially in their potential impact (Fig.1), even for those situated within the boundary of the ACEC (Fig. 2). The second case study demonstrated how to assess where anthropogenic noise is likely to be audible to humans, whereas the third case study examined the potential for noise to mask species communications.

These case studies provided insights useful for management and noise mitigation. The first case study identified the potential for alternative, quieter well locations, and highlighted which well sites would benefit from additional noise-control measures. In the second case study, mitigation actions could be taken for any locations where audible anthropogenic noise would be unacceptable. Further, the analysis highlighted which regions of the motorized trail were most responsible for the noise. These areas could be the focus of mitigation efforts (e.g., by re-routing trails). The results from the third case study could be used to develop restrictions on the number or types of snowmobiles allowed in an area or to identify particularly sensitive areas to be closed to recreational use.

The analyses required several modeling decisions, and the appropriate decision depended on the specific question being asked in each case study. These included the choice of acoustic metric, sound propagation model, source level data and number of sources, resolution and extent of the analysis, weather data and season, and alternatives for evaluation of planning and mitigation options. Modeling decisions may have important effects on sound level predictions. Where there is uncertainty regarding the correct choice, we recommend running model predictions for several input conditions to assess the sensitivity of the predictions to the modeling decisions. Consistent results will highlight robust model predictions, whereas divergent results will require careful consideration. Below, we examine the influence of modeling decisions in the three case studies.

These case studies highlighted that a variety of acoustic metrics are available, including sound pressure levels (Fig. 1a), thresholds (Fig. 1b), audibility (Fig. 3), and potential for masking (Fig. 4). Each of these metrics can give useful information, but regardless of the choice of metric, it is critical to report enough detail to allow the metric to be interpreted and converted to other metrics for comparison purposes (reviewed by McKenna, Shannon & Fristrup 2016). In the first case study, we chose a threshold-based metric to provide a discrete value that could be used for interpreting relative ecological impacts among well locations. A weakness of thresholds is that they can be somewhat arbitrary (e.g., a mule deer could respond similarly to values immediately below and above the selected threshold), but in this case study, the threshold approach provided a clear basis for comparing the relative impacts of the different well locations. In the second case study, we chose to examine audibility, as it may determine a minimum level of impact (i.e., audible or not). However, audibility will depend on species, individual, and even the degree of attention paid by an individual animal to the noise source (Fay 1988; Rapoza, Sudderth & Lewis 2015). Furthermore, a sound may be audible without necessarily causing any negative effects (Rapoza, Sudderth & Lewis 2015). Finally, in the third case study, we chose to examine the potential for masking, as acoustic communication is critically important to many species’ survival and reproduction. Anthropogenic noise can disrupt communication by masking signals or requiring additional energy expenditures to counteract the effects of the noise (Brumm & Slabbekoorn 2005).

Once the appropriate acoustic metric has been decided, it may be important to select a particular sound propagation model. The choice of sound propagation model can be guided by empirical data (e.g., see Appendix 4), consideration of the model capabilities (e.g. frequency range, land cover, see Keyel et al. *in review* for a detailed comparison of the three models in SMT), and applicable standards (e.g., Sunder 2003). This was exemplified in the third case study, where NMSIMGIS was chosen because it incorporated the necessary land cover types and frequency ranges.

Source type and source number can strongly influence the model results (Fig. 4). The propagation models described here provide an efficient means to examine a range of source types and numbers. Once the model has been run for a single source, sound levels from additional sources of the same type and at the same location can be quickly calculated, as sound levels increase by 3 dB per doubling of the number of sources (Bies & Hansen 2009). One approach to overcome the challenge presented by multiple sources of different types is to focus on total noise levels rather than numbers of vehicles, as was done in the winter travel management plan for Yellowstone National Park (Jacobson 2013). In the absence of specific source data, model predictions may be used to identify locations on the landscape more susceptible to noise intrusions. Finally, in particular management contexts, empirical data on the number, types, and use patterns of sources may be desirable.

Related to source number and type was the decision of how to represent line and area sources as points. In the second case study, we chose a spacing that gave sufficient coverage to examine relative impacts of different sections of the motorized trail. However, point sources differ from line sources in how fast sound levels drop (6 dB per doubling of distance for point sources compared to 3 dB for line sources, Bies & Hansen 2009). Consequently, where accurate representation of a line is required, it will be necessary to check that the chosen point spacing accurately models the 3 dB loss per doubling of distance. For the third case study of the snowmobile area, a coarser grid may be sufficient and is more computationally efficient for evaluating greater spatial extents.

Choice of weather conditions can also influence the model results. Here, we chose to use weather data from a nearby weather station using seasonally appropriate weather conditions. Our goal was not a precise instantaneous sound level for one given point in time, but a relative assessment of the different options under equivalent conditions (see Appendix 3 for a broader discussion of weather-related model considerations).

There were several options for considering alternative source placement. In the first case study, alternatives were examined with a systematic grid of points. In the second case study, alternative possible routes could have been considered using the same methods as were used for the existing motorized trail. A third option, not demonstrated here, would have been to use randomly placed points in order to provide spatially unbiased inferences (Quinn & Keough 2002 pp. 155 – 157). Finally, consideration of alternatives may benefit from additional information, for example by excluding impractical locations from the analysis.

Once the modeling decisions have been made, and the analyses completed, it is important to recognize the limitations of the results. For example, the first case study highlighted an important omission in our analysis: we considered only the noise impacts of the well site itself, and not those of any associated infrastructure (e.g., roads). A quiet well location that would require a noisy access road through sensitive areas may be worse than an alternative well location with quieter access. Further analyses would need to be carried out to examine the cumulative noise impacts of the well site decision. Finally, management decisions would need to consider information beyond just noise (e.g., presence of sensitive species, wilderness characteristics, access to the resource of interest, sensitivity of the habitat to disturbance, etc.).

Similarly, the second case study was limited by the use of estimated background levels that did not include other anthropogenic noise sources, such as a nearby highway. Further, while a single ATV may not be audible above background sound levels, a louder model or multiple ATVs may be audible for greater portions of the trail. An additional limitation for all three case studies was that sound sources were assumed to be omni-directional, although this may not be true for some sound sources (e.g., Conner & Page 2002).

The third case study could also be improved by incorporating additional factors in to the analysis to strengthen its application to management. Specifically, masking depends on signal amplitude (i.e. how loud a nuthatch vocalizes), the range of frequencies used for communication, how well a species can hear during noise events (e.g., through the use of critical ratios, Lohr, Wright & Dooling 2003), and whether the species behaviorally adjusts for the noise (e.g., by shifting its vocalization frequency or timing of vocalizations, Brumm & Slabbekoorn 2005; Slabbekoorn 2013). Quantitative estimates of masking could be made using a species’ signal amplitude for important vocalization frequencies in conjunction with its ability to hear those frequencies in noise (Lohr, Wright & Dooling 2003).

### SYNTHESIS AND APPLICATIONS

In these case studies, we demonstrated a robust modeling framework for evaluating the potential noise impacts likely to be encountered by animals in natural areas. This framework can be used to evaluate alternative management scenarios, such as to address changes to current management practices or in advance of decision-making for the siting of noise sources. This approach could be used to designate quiet areas, where any noise intrusion would be harmful to animal species, areas where noise impacts should be mitigated (e.g., through use of quieter technology, Bayne, Habib & Boutin 2008), and areas where additional noise sources are unlikely to provide an appreciable increase above background levels or negatively impact critical resources. This framework provides an approach to reduce the negative impacts of noise pollution and thereby promote the conservation of biodiversity.

Finally, all of the approaches employed here used a single sound level per noise source, while many noise sources vary in decibel level over time. More sophisticated examples could have been developed that used more than one source level (e.g. for multiple snowmobile speeds, multiple vehicle types, or for a source that varies in sound level over time), and summary information could have been extracted such as maximum sound level, duration of the noise source, time audible, and whether the noise source is impulsive (such as a gunshot) or continuous. However, more studies are needed to understand how these different metrics relate to animals’ behavior and fitness (Shannon *et al*. 2015) and we believe the modeling tools utilized here can facilitate such studies for terrestrial species.

## Author Contributions

ACK, SER, and KN conceived the ideas and designed methodology; ECM, JB and ACK collected the data; ACK analyzed the data; ACK led the writing of the manuscript, and GW, SER, and KN revised it critically for important intellectual content. All authors contributed critically to the drafts and gave final approval for publication.

## Acknowledgements

We thank J. Slivka, P. Hartger, and The Wilderness Society for suggesting the case studies and providing data, E. Brown for the drill rig data, D. Joyce for the snowmobile data, and J. Thomson, the Reed Lab and the National Park Service Natural Sounds and Night Skies Division for constructive feedback and discussions. This research was funded by a grant to the Wildlife Conservation Society from the William and Flora Hewlett Foundation and the National Park Service.

## Data Accessibility

–Sound Mapping Toolbox is available from: http://purl.oclc.org/soundmappingtools
–Source level data references given in Table 1
–Python scripts: uploaded as online supporting information
–Public data sets: Area of Environmental Concern (BLM 2015b), BLM trail designations (Zone P, BLM 2015c), roads (U.S. Census Bureau 2016), Elevation Data: (USGS 2013), Land cover data: (LANDFIRE 2012), Weather data: See (NOAA 2015), aerial photos (USDA-FSA-APFO 2016), White-breasted Nuthatch recordings (Nelson 2015a; b)

## Supporting Information

**Appendix 1**: The scripts used to run the examples

**Appendix 2:** Calculation of mule deer threshold

**Appendix 3:** Weather-related considerations

**Appendix 4:** Use of empirical data to guide sound propagation model selection

